# The chaperonin CCT8 controls proteostasis essential for T cell maturation, selection, and function

**DOI:** 10.1101/2020.08.11.246215

**Authors:** Bergithe E Oftedal, Stefano Maio, Adam Handel, Madeleine PJ White, Duncan Howie, Simon Davis, Nicolas Prevot, Ioanna A. Rota, Mary E Deadman, Benedikt M Kessler, Roman Fischer, Nikolaus S Trede, Erdinc Sezgin, Rick M Maizels, Georg A Holländer

**Affiliations:** Developmental Immunology, MRC Weatherall Institute of Molecular Medicine, University of Oxford, Oxford OX3 9DS, United Kingdom; Department of Clinical Science, University of Bergen, Bergen, Norway; K.G. Jebsen Center for Autoimmune Disorders, Bergen, Norway; Wellcome Centre for Integrative Parasitology, Institute of Infection, Immunity and Inflammation, University of Glasgow, G12 8TA, United Kingdom; Sir William Dunn School of Pathology, University of Oxford, Oxford, OX1 3RE, United Kingdom; Target Discovery Institute, Nuffield Department of Medicine, University of Oxford, Old Road Campus, Headington, Oxford, OX3 7FZ, United Kingdom; Huntsman Cancer Institute, University of Utah, Salt Lake City, UT, USA; MRC Human Immunology Unit, MRC Weatherall Institute of Molecular Medicine, University of Oxford, Oxford OX3 9DS, United Kingdom; Paediatric Immunology, Department of Biomedicine, University of Basel, Basel, Switzerland; Department of Biosystems Science and Engineering, ETH Zurich, Basel, Switzerland

**Keywords:** Chaperonin CCT8, T cell biology, nuclear actin, helminth infection

## Abstract

T cells rely for their development and function on the correct folding and turnover of proteins generated in response to a broad range of molecular cues. In the absence of the eukaryotic type II chaperonin complex, CCT, T cell activation induced changes in the proteome are compromised including the formation of nuclear actin filaments and the formation of a normal cell stress response. Consequently, thymocyte maturation and selection, and T cell homeostatic maintenance and receptor-mediated activation are severely impaired. Additionally, Th2 polarization digresses in the absence of CCT-controlled protein folding resulting paradoxically in continued IFN-γ expression. As a result, CCT-deficient T cells fail to generate an efficient immune protection against helminths as they are unable to sustain a coordinated recruitment of the innate and adaptive immune systems. These findings thus demonstrate that normal T cell biology is critically dependent on CCT-controlled proteostasis and that its absence is incompatible with protective immunity.

## Introduction

T cells are indispensable co-ordinators of the adaptive immune response and embrace different effector functions dependent on the context in which they recognize their cognate antigen (Yamane & Paul, 2012). The competence of T cells to respond adequately to antigenic challenges is inextricably linked to *de novo* protein expression and a change in the cell’s protein homeostasis. Under optimal conditions this process relies on a complex network of interconnected, dynamic systems that control protein biosynthesis, folding, translocation, assembly, disassembly, and clearance (Balch, Morimoto et al., 2008). Cell stress and other physiological demands on the cell’s proteome can however result in challenges where nascent and metastable proteins are misfolded and, as a result, aggregate–entrapped polypeptides are formed. These conformational changes challenge, as toxic intermediates, a cell’s functions and may impair its survival. Thus, protein quality control and the maintenance of proteostasis are essential for almost all biological processes. This is in part accomplished by a machinery of chaperones that catalytically resolve misfolded proteins from adopting a state of amorphous aggregates but assist them in assuming a native conformation (Cao, Carlesso et al., 2008, Morimoto, 2008).

Chaperones are operationally defined by their capacity to interact transitorily with other proteins in assisting *de novo* folding of nascent proteins, refolding of stress-denatured proteins, oligomeric assembly, protein trafficking and proteolytic degradation (Hartl & Hayer-Hartl, 2009). Chaperones that partake in *de novo* folding or refolding promote these conformational changes through the recognition of hydrophobic amino acid side chains of non-native polypeptides and proteins. Eukaryotic Type II chaperones, aka chaperonins, are large (~800−900 kDa), cytoplasmic, protein complexes (designated CCT, <u>chaperonin containing tailless complex polypeptide 1, TCP-1 or TCP-1-ring complex, TRiC</u>), that are built as hetero-octameric cylinders formed from two stacked doughnut-like rings (Leitner, Joachimiak et al., 2012). Each of the rings is composed of homologous, yet distinct 60 kDa subunits (α, β, γ, δ, ε, ζ, η and θ (Spiess, Miller et al., 2006). CCTs recognize, bind and globally enclose protein substrates of up to ~60 kDa to allow their folding over several cycles. Multiple substrates are recognized by CCTs whereby each of the complex’s subunits may identify different polar and hydrophobic motifs (Joachimiak, Walzthoeni et al., 2014). High affinity substrates are evicted from CCTs in an ATP-dependent fashion as they act as competitive inhibitors of the complex’s catalytic reaction of unfolding proteins (Priya, Sharma et al., 2013).

As many as 10% of newly synthesized proteins are assisted by CCT to adopt a correct conformation (Yam, Xia et al., 2008), including key regulators of cell growth and differentiation, and components of the cytoskeleton (Dekker, Stirling et al., 2008, Grantham, Brackley et al., 2006). Actins and tubulins have been identified as two of the major folding substrates of CCT (Sternlicht, Farr et al., 1993). Adequate production of effector cytokines by T cells has been related to rapid actin polymerization and the generation of a dynamic filament network in the nucleus of CD4^+^ T cells (Tsopoulidis, Kaw et al., 2019). Moreover, signal transduction, cytoskeletal synthesis and remodelling, immune synapse formation, macromolecular transport, and cell division require molecules that depend on CCT function (Dustin & Cooper, 2000, Gunzer, Schafer et al., 2000).

To dissect the precise role of CCT in T cell development and function we generated mice that lack the expression of a single subunit, CCTθ(a.k.a. CCT8), in immature thymocytes and their progeny. We tested the proficiency of these cells to normally develop, be selected within the thymus, and respond to antigens as part of an adaptive immune response. Our results show that CCT8 is largely, albeit not entirely, dispensable for thymocyte differentiation and selection but essential for mature T cells to respond adequately to antigenic stimuli.

## Results

### Thymocyte development and peripheral T cell differentiation depend on CCT8 expression

The subunit 8 of Type II chaperones, CCT8, was detected throughout thymocyte development but was most prominently found in immature cells with a double negative (DN, i.e. CD4^−^CD8^−^) phenotype (**Figure 1a**). CD4-Cre::CCT8^fl/fl^ mice (designated CCT8^T−/−^; **Figure EV 1a**) have a loss of CCT8 expression targeted to double positive (DP i.e. CD4^+^ and CD8^+^) thymocytes and their progeny (**Figure EV 1b, Appendix figure 1a**). The total thymus cellularity of these mice was comparable to that of Cre-negative littermates (designated CCT8^T+/+^; **Figure 1bi**). As expected, the frequency of DN and immature single positive (iSPCD8) thymocytes remained unaffected as the deletion of CCT8 occurs after these maturational stages (**Figure 1bii, Figure EV 1c**). However, the frequency of DP thymocytes was mildly increased and their progression to a post-signalling stage (TCRβ^hi^CD69^−^) was impaired (**Figure 1c, Appendix figure 1b**), which correlated with a higher number of thymocytes that had not received a sufficiently strong survival signal (**Figure 1d, Appendix figure 2**). In parallel, the extent of negative selection during the first selection wave, as identified by the co-expression of Helios and PD1 on Foxp3^−^CCR7^−^TCR^+^ DP or CD4 thymocytes (a.k.a. wave 1a and b, respectively), was reduced in CCT8^T−/−^mice (**Figure 1e, Appendix figure 3**). The following wave, which takes place in the medulla and which is characterized by Helios expression on Foxp3^−^ SPCD4 thymocytes, was reduced in a first (CD24^+^) but not a second phase (CD24^−^; **Figure 1f**). Finally, fewer phenotypically and functionally mature single positive CD4^+^ (SP4) and SPCD8 thymocytes were detected at a late stage of their development (**Figures 1bii, and 1g**, **Figures EV 1d** and **1e**) and the frequencies of thymic and recirculating regulatory T cells (T_reg_) were likewise reduced (**Figure 1h, Appendix figure 4**). Hence, the targeted loss of CCT8 expression in thymocytes impaired their selection and reduced the frequency of post-selection, mature effector, and regulatory T cells.

**Figure 1.**
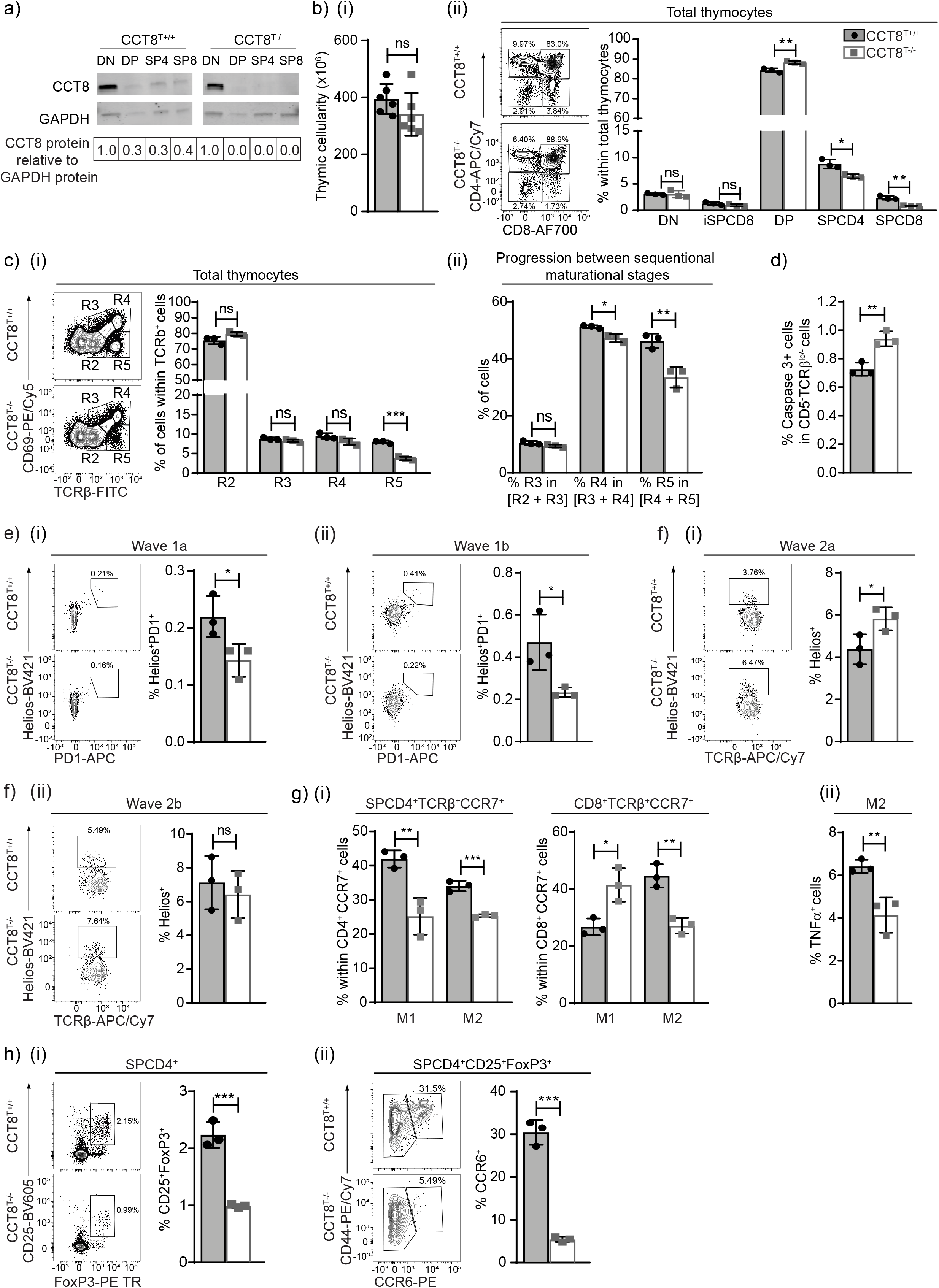
The role of CCT8 in thymocytes. (**a**) CCT8 protein detection by Western blot. Values shown are relative to the detection in DN thymocytes. (**b**) (i) Total thymic cellularity and (ii) thymocyte subpopulations in 6 week old CCT8^T+/+^ (grey bars) and CCT8^T−/−^ mice (white bars) as defined by CD4 and CD8 cell surface expression on lineage-negative thymocytes. (**c**) Positive thymocyte selection. (i) Maturational stages (R2: TCRβ^low^CD69^−^; R3: TCRβ^low^CD69^+^, R4: TCRβ^hi^CD69^+^, and R5: TCRβ^hi^CD69^−^) and (ii) progression between sequential stages. (**d**) Non-signaled thymocytes (activated caspase 3- expression among CD5^−^TCRβ^lo^/^−^ thymocytes). (**e**) Negative thymocyte selection in the cortex. (i) Wave 1a (TCRβ^lo/hi^ CCR7^−^ SPCD4^+^SPCD8^+^) and (ii) wave 1b (TCRβ^hi^ SPCD4^+^CCR7^−^CD69^+^CD24^+^). (**f**) Negative selection in the medulla. (i) Wave 2a (TCRβ^hi^SPCD4^+^CCR7^+^CD69^+^CD24^+^) and (ii) wave 2b (TCRβ^hi^SPCD4^+^CCR7^+^CD69^−^CD24^−^). (**g**) Late stage maturation of single positive TCRβ^hi^CCR7^+^ thymocytes. (i) Phenotypic analysis (M1: CD69^+^MHC^+^, M2: CD69^−^MHCI^+^). (ii) TNFα production by M2 SPCD4. (**h**) (i) Total thymic T_reg_ cells (CD25^+^Foxp3^+^), and (ii) recirculating T_reg_ among thymic CD25^+^Foxp3^+^ cells. Left panels in bii, ci, ei, eii, fi, fii, hi, and hii, display representative contour plots of data shown in bar graphs (b-h). *p<0.05, **p<0.01, ***p<0.001, ****p<0.0001 (Student’s t test, b-h, adjusted for multiple comparison panel gi). Data shown in panel (a) are representative of 2 independent experiments. Data in bar graphs show the mean ±SD representative two (panels b-d, and g) and three (e, f, h) independent experiments, respectively with each three replicates each. See also Figure S1 and Appendix figure 1-4.

The total splenic cellularity of CCT8^T−/−^ mice was normal, although drastically fewer naïve T cells were detected (**Figure 2a and 2b**, **Appendix figure 5a**) and the CD4 and CD8 lineages were differentially affected (**Figure 2c**). The frequencies of CD4 and CD8 T cells with a memory phenotype was increased (**Figure 2d**), likely reflecting homeostatic expansion as a consequence of low T cellularity. Correspondingly, the frequency of peripheral T_reg_ cells (CD25^+^FoxP3^+^) remained unaffected in CCT8^T−/−^ mice but the subpopulation of highly suppressive CD103^+^ ICOS^+^ T_reg_ was several fold increased in line with the extent of lymphopenia (**Figure 2e, Appendix figure 5b)** (Barthlott, Bosch et al., 2015). Moreover, the frequency of CD4^+^ memory T cells with an anergic phenotype (CD73^+^FR4^+^) was reduced in CCT8^T−/−^ mice (**Figure 2f, Appendix figure 5a**). Collectively and contrary to the relatively minor decrease in thymic SP cells, the loss of CCT8 expression correlated with a severe reduction in peripheral T cells implying a functional impairment of these cells.

**Figure 2.**
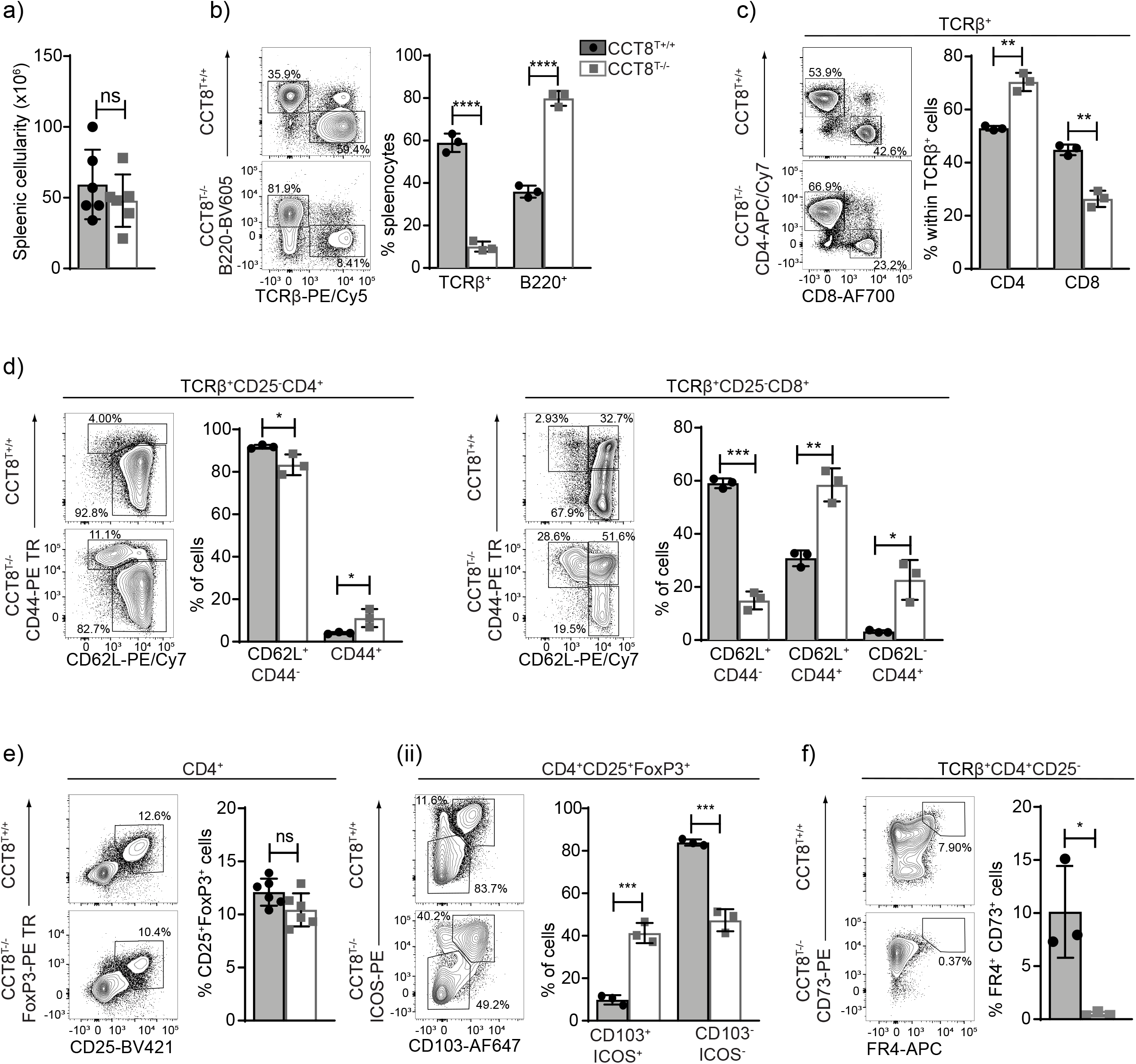
Peripheral T cell cellularity and phenotype in the absence of CCT8 expression. Analysis of 4-6 week old CCT8^T+/+^ (grey bars) and CCT8^T−/−^ mice (white bars). Gating strategies are shown for representative contour plots. (**a**) Total splenic cellularity. (**b**) Frequencies of splenic B and T cells. Frequencies **(c)** of total splenic CD4 and CD8 T cells, and (**d**) their naïve (CD62L^+^CD44^−^) and memory, effector memory (CD62L^−^CD44^+^), and central memory subpopulations (CD62L^+^CD44^+^), respectively. (**e**) The frequencies of (i) total splenic T_reg_ cells, and (ii) T_reg_ cell subpopulations defined by ICOS and CD103 cell surface expression. **f**) Frequency of anergic CD4^+^ T cells in lymph nodes. *p<0.05, **p<0.01, ***p<0.001, ****p<0.0001 (Student’s t test, a-f). Contour plots (b-f) are representative of data in bar graphs. Data shown in bar graphs represent mean ±SD values of a single experiment and are illustrative of two independent experiments with three replicates each. See also Appendix figure 5.

### Loss of CCT8 impairs the formation of nuclear actin filaments

Activated naïve CCT8^T−/−^ T cells displayed a drastically reduced expansion index when compared to controls and their survival was greatly reduced, which was enhanced by neither the addition of IL-2 nor anti-oxidant N-Acetyl-L-cysteine (**Figure 3a** and **Figure EV 2a**) (Eylar, Rivera-Quinones et al., 1993). Because T cell activation is associated with significant *de novo* protein synthesis (Brownlie & Zamoyska, 2013) and the engineered lack of CCT8 expression reduced the expression of all components of the CCT complex (**Figure 3bi**), we next quantified in CCT8-deficient and - proficient T cells changes in tubulin and actin expression, as these two serve as folding substrates for the CCT complex (Grantham et al., 2006, Saegusa, Sato et al., 2014). Several tubulin isoforms were reduced in CCT8^T−/−^ T cells, independent of the activation state, while the detection of the ubiquitously expressed β and γ actin isoforms was unaffected by a loss of normal CCT expression (**Figure 3bii**). However, the formation of nuclear actin filaments could only be detected in a very small fraction of CCT8-deficient T cells (3.17 ± 1.15 % versus 29.40 ± 10.5) similar to the frequency of wild type T cells in which Arp2/3 was pharmacologically inhibited and actin nucleation was prevented (**Figure 3c**) (Thiam, Vargas et al., 2016). Hence, the formation of nuclear actin filaments, an important requirement for T cell function (Tsopoulidis et al., 2019), was dependent on an intact CCT complex.

**Figure 3.**
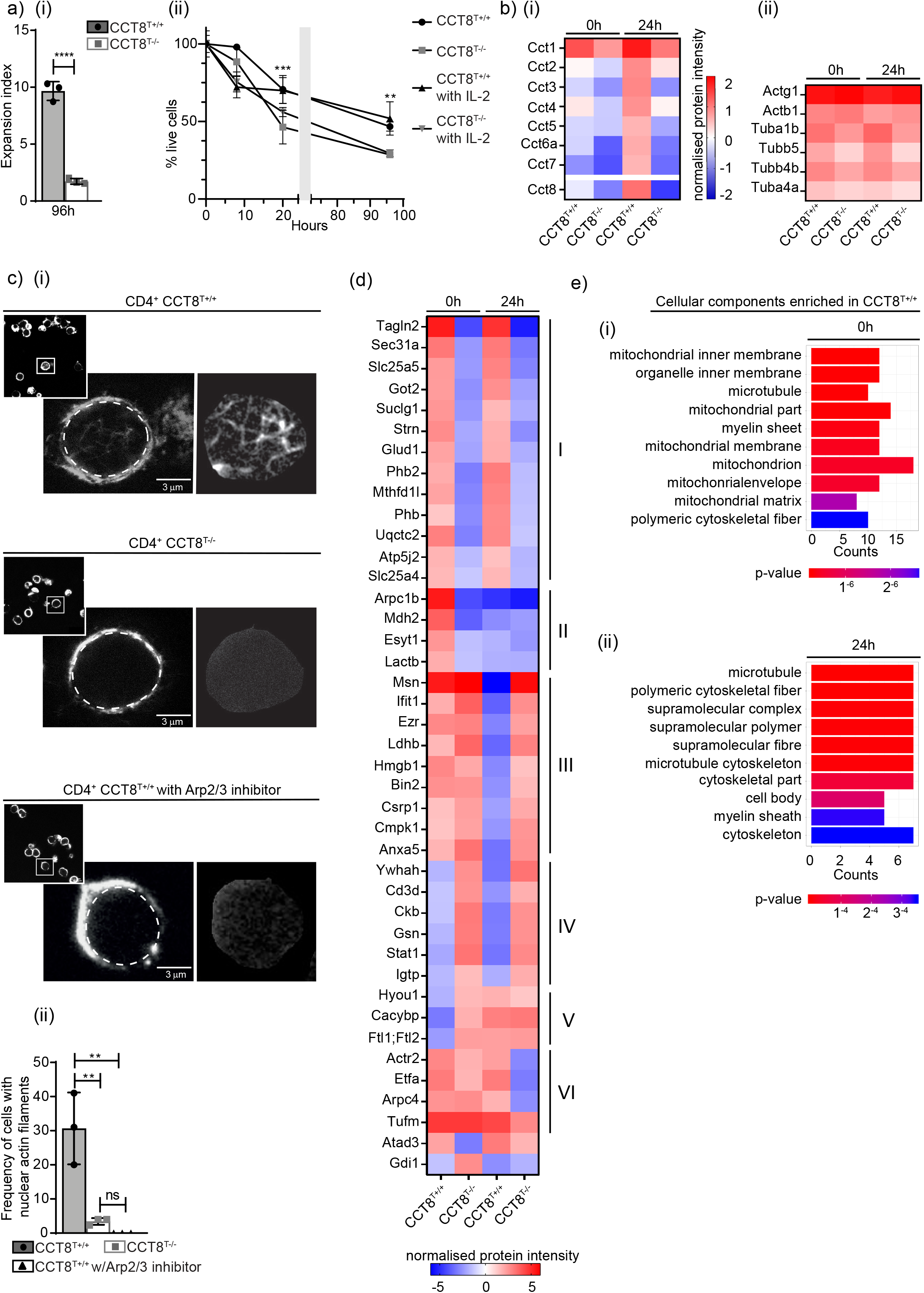
Proteomic analysis of resting and activated CD4^+^ T cells from CCT8^T+/+^ and CCT8^T−/−^ mice. (**a**) T cells activated *in vitro by* CD3/CD28 cross linking. (i) Expansion index; (ii) normalised cell viability in the presence or absence of exogenous IL-2. (**b**) Expression of individual CCT subunits (i) and the CCT substrates actin and tubulin measured by mass spectrometry, (**c**) Nuclear actin filaments in activated CD4^+^ T cells in the presence of absence of CK-666, an inhibitor of Arp2/3. (i) STED images with segmentation of the nuclear actin filaments (right panels), and (ii) frequency of nuclear actin filaments detected in CCT8^T+/+^ (white bars) and CCT8^T−/−^ T cells (grey bars). (**d**) Differentially expressed proteins in resting (0h) and activated (24h) T cells. (**e**) Go analysis of cellular components enriched in CCT8^T+/+^ CD4^+^ cells at 0 hrs and 24 hrs (ii) after stimulation **p<0.01, ***p<0.001, ****p<0.0001. Data was calculated by Student’s t test (ai and c) correcting for multiple comparisons (Holm-Sidak method; aii) and Anova with post hoc test (b and d). Data shown in bar graphs represent mean ±SD values of a single experiment representative of two independent experiments with three replicates each, and results in heat maps and GO analyses are from two independent experiments with three replicates each. See also Figure S2 and Appendix figure 6.

### The lack of CCT8 compromises proteostasis in both resting and activated T cells

Upon T cell activation, the lack of CCT8 also changed the expression of proteins other than individual CCT components and tubulin (**Figure 3d**). At least six different patterns of protein changes were detected when comparing mutant and wild type T cells before and 24 hours after activation, including proteins that were minimally expressed in both resting and activated CCT8^T−/−^ T cells (groups I+II), or that could be detected in un-stimulated CCT8^T+/+^ T cells but following activation did not change (groups III-V) or even a decrease in expression was detected (VI) (**Figure 3d**). Many of the group I proteins were related to mitochondrial functions, for example the ATP synthase subunit beta (Atp5b, catalysing ATP synthesis (Prescott, Bush et al., 1994)), glutamate dehydrogenase 1 (Glud1, catalysing the oxidative deamination of glutamate (Karaca, Frigerio et al., 2011)), and prohibitin (PHB, controlling mitochondrial biogenesis but also cell-cycle progression, nuclear transcription and resistance to various apoptotic stimuli) (Merkwirth, Dargazanli et al., 2008, Peng, Chen et al., 2015). CCT8^T+/+^ CD4 T cells showed increased expression of mitochondrial-associated proteins and genes relative to CCT8^T−/−^ CD4 T cells, which was in agreement with the cells’ inadequate ability to increase protein synthesis and to adapt to metabolic demands (**Figure 3e** and **Figure EV 2b**). This deficiency in mitochondrial-related pathways was a consistent finding in both the proteome and transcriptome (**Figure 3e** and **Figure EV 2b**), and particularly focused on an altered expression of coenzyme Q10 metabolism and ATP synthesis. However, neither the biogenesis nor the membrane potential of mitochondria were impaired in activated T cells (**Figure EV 2c-e, Appendix figure 6a and b**). We noticed in resting T cells a reduction in the concentration of Arpc1b (group II) and a lack of an up-regulation of Arpc4 (group VI), in activated T cells, representing two of the five essential and non-interchangeable components of the Arp2/3 complex which promotes filamentous (F) actin branching (see **Figure 3c**) (Goley & Welch, 2006). The organizer proteins Moesin (Msn) and Ezrin (Ezr) (Group III) that link F-actin to the plasma membrane, remained highly expressed in stimulated CCT8^T−/−^ T cells thus impairing the remodeling of the cytoskeleton upon activation (Garcia-Ortiz & Serrador, 2020). Furthermore, several proteins failed to be reduced in response to CD4^+^ T cell activation (group III) including Ifit1, a protein expressed in response to interferons and the subsequent recruitment of STAT1 that also negatively regulates pro-inflammatory genes (John, Sun et al., 2018). RNA-Seq analysis showed differential gene expression and confirmed for activated CCT8^T−/−^ T cells an enrichment of genes belonging to the IFN-γ pathway (GO:0034341; 8.8-fold change in mutant when compared to wild type cells, adjusted p-value = 0.0002; **Figure EV 2f,g**) (Ramana, Gil et al., 2002).

### CCT8 is essential to avert T cell activation-induced cellular stress

The accumulation of unfolded protein in the endoplasmatic reticulum (ER) leads to cellular stress and, where unresolved, a loss of regular cell functions prompting apoptosis. Eukaryotic cells have developed an evolutionary well conserved mechanism to clear unfolded proteins and to restore ER homeostasis, known as the unfolded protein response (UPR) (Kemp & Poe, 2019). The UPR comprises a tightly orchestrated collection of signaling events that are controlled by protein kinase RNA-like ER kinase (PERK), activating transcription factor 6 (ATF6) and inositol requiring protein-1α (IRE-1α), which collectively sense ER stress and alleviate the accumulation of misfolded proteins, for example via increasing the expression of ER chaperones (Sano & Reed, 2013) (**Figure EV 3a**).

Increased transcripts for *Perk*, *Atf6* and *Ire1-α* were detected in activated CCT8^T−/−^ T cells when compared to controls (**Figure 4a**). In keeping with and consequent to an activated UPR, transcripts were increased for the chaperone glucose-regulated protein (Grp) 78, a target gene of ATF6, and XBP-1 which is placed in the ER lumen and contributes there to ER homeostasis, protein folding and degradation (Lee, Iwakoshi et al., 2003, Sano & Reed, 2013). Activation of IRE1-α also led to an up-regulation of pro-apotoptic *Bim* and a decreased expression of the anti-apoptotic *Bcl2*, thus contributing to the impaired survival of activated CCT8^T−/−^ T cells (**Figure 2a**). Taken together, activated CCT8^T−/−^ T cells displayed an extensive UPR as a result of impaired proteostasis, which appeared incompatible with normal T cell function.

**Figure 4.**
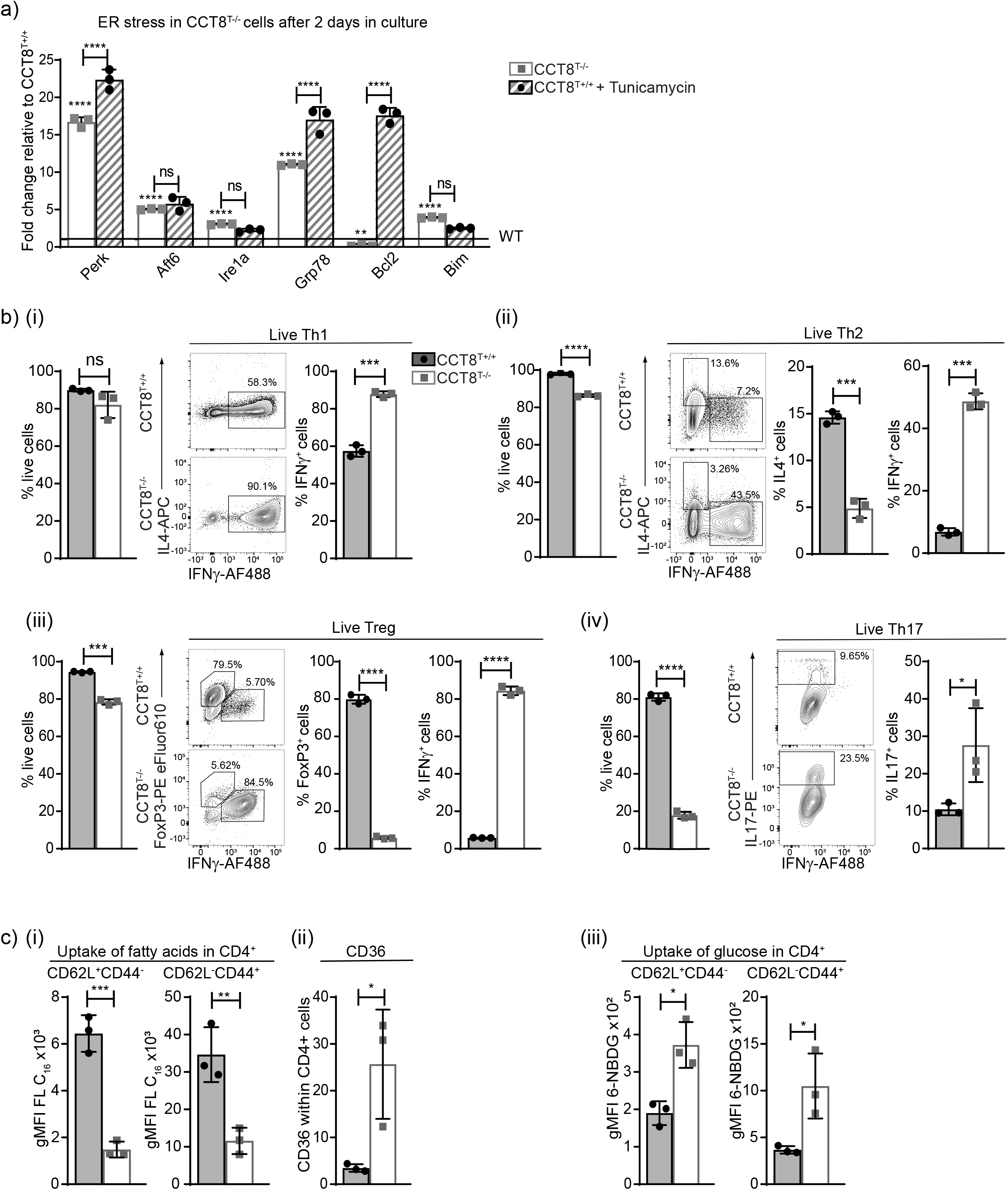
Peripheral T cell functions in the absence of CCT8 expression. Analysis of naïve CD4^+^ T cells from 4-6 week old CCT8^T+/+^ (grey bars) and CCT8^T−/−^ mice (white bars). (**a)** qPCR analysis of ER stress response elements in T cells activated by CD3 and CD28 crosslinking and cultured in the presence or absence of tunicamycin; expression normalised to GPDH and displayed as 2^−ΔΔCt^ CT values relative to values from CCT8^T+/+^ T cells arbitrarily set 1. (**b**) *In vitro* differentiation of peripheral naïve CD4^+^ T cells grown for 5 days under differentiating conditions: frequency of live cells (left) and cells (right) adopting (i) Th1 polarisation; (ii) Th2 polarisation (iii); T_reg_ differentiation; and (iv) Th17 differentiation. (**c**) (i) Uptake of fatty acids in CD4+ CD62L+CD44-(naïve) and CD62L-CD44+ (memory) cells *ex vivo* activated by CD3 and CD28 cross-linking for 20h; (ii) expression of long-chain fatty acid receptor CD36 on CD3/CD28-activated cells, and (iii) uptake of glucose analogue 6-NBDG in CD4+ CD62L+CD44-(naïve) and CD62L-CD44+ (memory) cells.*p<0.05, **p<0.01, ***p<0.001, ****p<0.0001. Data was calculated by Student’s t-test (panel a-c) and adjusted for multiple comparisons (panel a). Bar graphs show the mean ±SD and are representative for two independent experiments with three replicates each. See also Figure S3 and Appendix figure 7.

### Th2 cell polarization and T cell metabolism are dependent on CCT8 expression

Activated T cells undergo clonal expansion and differentiate, in the presence of additional molecular cues, into functionally distinct T subsets characterized by separate cytokine profiles and effector behaviours. Because mitochondrial and proteomic reprogramming parallel this differentiation we next examined whether an inadequate mitochondrial response to metabolic demands, combined with an abnormal UPR (**Figure EV 3a**), could impair peripheral T cell differentiation. Under Th1 polarizing conditions, the absence of CCT8 expression did not impair the viability of *in vitro* activated CD4 T cells but increased their frequency to express the signature cytokine IFN-γ (**Figure 4bi, Appendix figure 7a**). In contrast, both viability and IL-4 production were reduced in activated CD4 CCT8^T−/−^ T cells polarized to adopt a Th2 phenotype (**Figure 4bii, Appendix figure 7b**). Paradoxically, the frequency of IFN-γ expressing CD4 CCT8^T−/−^ T cells was increased by 7-fold under these conditions. Both of these finding are in agreement with an increased detection of STAT1 in CCT8^T−/−^ T cells, independent of their activation status (**Figure 3d**). Conditions favouring T_reg_ differentiation not only caused a reduced viability in CCT8^T−/−^ T cells but also resulted in a reduced conversion of these cells to express FoxP3, whereas the vast majority of viable cells unexpectedly expressed IFN-γ (**Figure 4biii, Appendix figure 7c**). Driving CCT8^T−/−^ T cells to a Th17 phenotype correlated with a dramatic loss in viability albeit the frequency (but not the total cellularity) of living cells successfully polarized was increased (**Figure 4biv, Appendix figure 7d**). This reaction was CCT8^T−/−^ T cell-intrinsic, as a comparable response was observed when CCT8-deficient and -proficient T cells were co-cultured during polarization (**Figure EV 3b**). Thus, polarization of T cells was severely altered in the absence of CCT8, thus favouring these cells to adopt a Th1 phenotype.

As T cells transform from a quiescent to an activated state, the generation of energy from shared fuel inputs such as fatty acids and glucose are essential for the cells’ growth, differentiation and survival (Howie, Ten Bokum et al., 2017, O’Neill, Kishton et al., 2016). We therefore tested whether an absence of CCT8 expression impaired the use of these two essential energy sources by CD4^+^ T cells. Fatty acid uptake was significantly reduced in both activated naïve and memory CD4^+^ T cells lacking CCT8 (**Figure 4di**). This result was notably independent of an increased cell surface expression of CD36, a glycoprotein that acts together with chaperones to translocate fatty acids to the cytoplasm (**Figure 4dii**). In contrast to wild type controls, both naïve and memory CCT8^T−/−^ T cells displayed a higher glucose uptake (**Figure 4diii**) indicating a compensatory mechanism to be in play that secures, at least in part, the cells’ energy expenditure. Indeed, basal respiration, spare respiratory capacity, and extracellular acidification rate (ECAR), which measures glycolysis, remained globally normal despite a lack in CCT8 (**Figure EV 3c**). Hence, the absence of CCT8 in T cells resulted in a change in energy usage which may further explain the impaired response to cell activation.

### CCT8 is essential for protective immunity against intestinal helminths

Infection with the nematode *Heligmosomoides polygyrus* activates a strong Th2-type immune response (Maizels, Hewitson et al., 2012a). Primary infections in susceptible strains such as C57BL/6 are normally non-resolving and clearance requires drug treatment with pyrantel embonate. In contrast, spontaneous clearance of secondary infections relies on a protective anti-helminthic immunity and is observed in mice previously exposed to *H. polygyrus* and subsequently drug treated (Anthony, Urban et al., 2006). We therefore investigated whether the Th2 polarization deficiency observed in CCT8^T−/−^ mice would impair the expulsion of *H. polygyrus* after a short primary infection followed by a re-exposure to the parasite’s larvae. In comparison to wild type animals, a higher burden of worm eggs was detected in the faeces of infected CCT8^T−/−^ mice, especially on re-exposure (**Figure 5a**). This result correlated at the peak of primary and secondary infections with both fewer total cells and a reduced frequency of CD4^+^ Th2 cells (as determined by IL-4 and Gata3 expression) in mesenteric lymph nodes and peritoneal lavage (**Figures 5b,c, Appendix figure 8a**).

**Figure 5.**
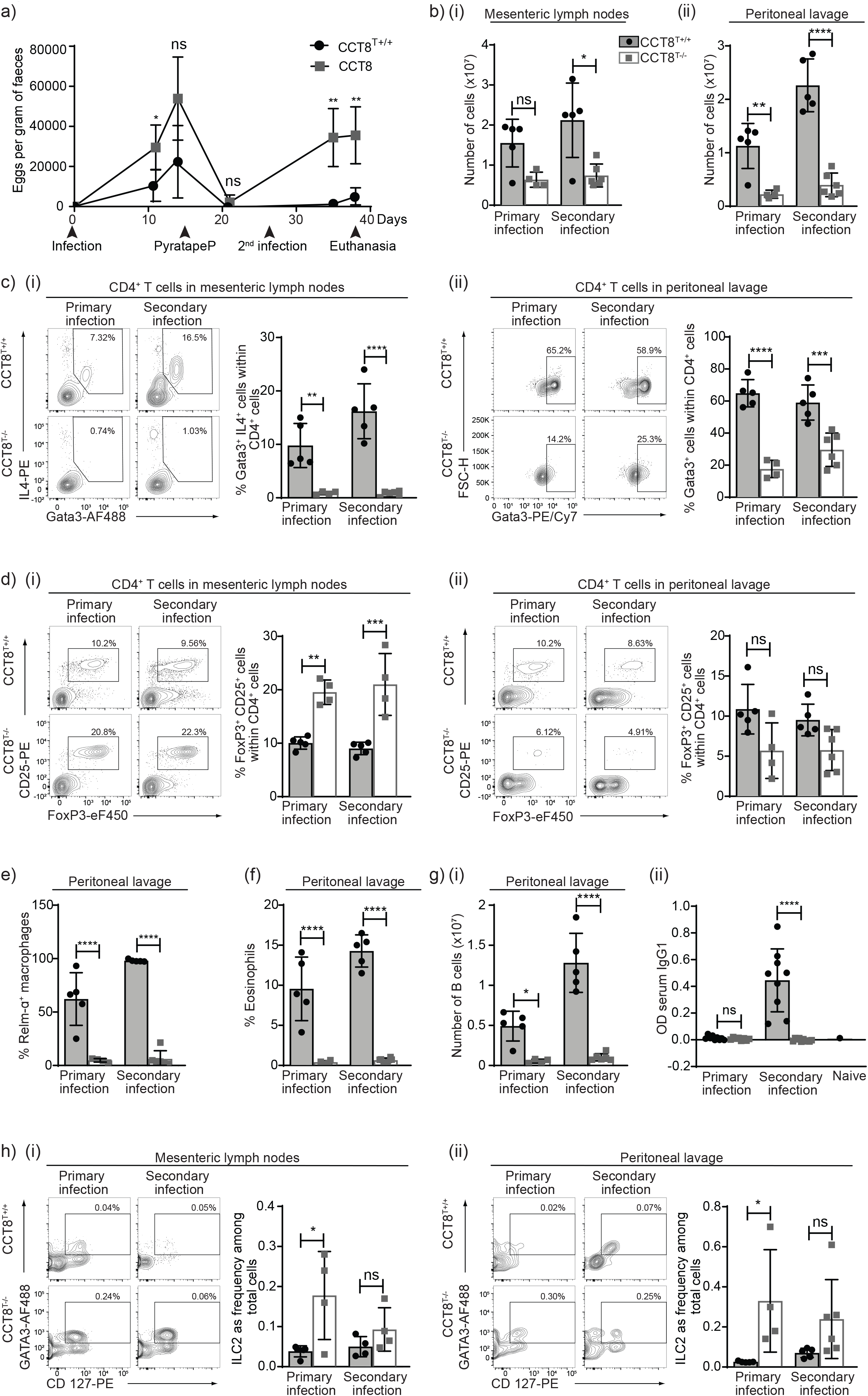
The response to primary and secondary *H. polygyrus* infection. Comparison of CCT8^T+/+^ (black circles) and CCT8^T−/−^ mice (grey squares). (**a)** Eggs per gram of faeces at indicated time points. (**b**) Cellularity in (i) mesenteric lymph nodes and (ii) peritoneal lavage. (**c**) Th2 cell frequency in (i) mesenteric lymph nodes and (ii) peritoneal lavage.(**d**) Treg frequencies in (i) mesenteric lymph nodes and (ii) peritoneal lavage. Frequencies of (**e**) alternatively activated macrophage and (**f**) eosinophils in peritoneal lavage. (**g**) (i) B cells in peritoneal lavage and (i) *H. polygyrus*-specific serum IgG1. (**h**) ILC2 frequency in (i) mesenteric lymph nodes and (ii) peritoneal lavage. Panels c, d, and h display contour plots with gating. *p<0.05, **p<0.01, ***p<0.001, ****p<0.0001. Data was calculated by Student’s t test adjusted for multiple comparison. Bar graphs show the mean ±SD and are representative of one of 2 independent experiments with at least 4 samples per group.

Resistance to *H. polygyrus* infections is closely related to the extent and speed of a pathogen-specific Th2 response whereas susceptibility to these worms is linked to T_reg_ activity and IFN-γ production (Sawant, Gravano et al., 2014, Smith, Filbey et al., 2016). We therefore analysed the frequency of T_reg_ populations during primary and secondary *H. polygyrus* infections. In mesenteric lymph nodes, CCT8-deficient T_reg_ cells were detected at higher frequencies during the primary and secondary pathogen challenges when compared to wild type controls (**Figure 5di, Appendix figure 8a**). However, fewer T_reg_ were observed in the peritoneal lavage of infected CCT8^−/−^ mice, possibly reflecting differences in their homing to or survival *in situ*, although these differences did not reach statistical significance (**Figure 5dii**). Hence, the insufficient priming and the subsequent absence of a protective immune response to a second *H. polygyrus* challenge correlated with the detected preference of CCT8^−/−^ T cells to adopt a Th17 phenotype, their readiness to secrete IFN-γ different activating and their reduced capacity to adopt a Th2 phenotype and paralleled a local expansion of T_reg_ cells in mesenteric lymph nodes in response to priming to *H. polygyrus*.

The exposure to Th2 cytokines in the context of parasite infections promotes granulomatous reactions composed of alternatively activated macrophages (AAMacs). Identified by their elevated expression of the resistin-like molecule (RELM)-α (Ariyaratne & Finney, 2019), AAMacs were recovered from the peritoneal lavage of wild type animals infected with *H. polygyrus* but were hardly detected at these sites in CCT8^T−/−^ mice, even after re-exposure to the parasite (**Figure 5e, Appendix figure 8b**). Similarly, the number of eosinophils in the peritoneal lavage was low in CCT8^T−/−^ mice with a primary *H. polygyrus* infection and this paucity remained unchanged upon re-infection (Figure 5f), thus correlating with the restricted capacity of CCT8^T−/−^ mice to be polarized to a Th2 phenotype.

*H. polygyrus* infections activate B cells and elicit a strong humoral immune response supporting the clearance of the parasite (Harris & Gause, 2011). We therefore also determined the absolute number and frequency of B cells in the peritoneal lavage of infected wild type and CCT8^−/−^ mice (**Figure 5gi, Appendix figure 8c**) and probed the secretion of *H. polygyrus*-specific serum IgG1. In comparison to infected controls, there was a higher frequency of B cells in the mesenteric lymph nodes of mutant mice but fewer in the peritoneal cavity during both primary and secondary infections (**Figure 5gi,**). In parallel, CCT8^T−/−^ mice failed to generate an antigen-specific IgG1 response to a secondary *H. polygyrus* infection (**Figure 5gii**) further highlighting the consequences of a limited Th2 response *in vivo*.

The type 2 cytokines IL-5 and IL-13 are also secreted early during a helminth infection by innate lymphocytes (ILC2) in response to the release of IL-33 and TSLP from mucosal epithelial sensor cells (Cortes, Munoz-Antoli et al., 2017, von Moltke, Ji et al., 2016). We therefore quantified these innate lymphocytes in mesenteric lymph nodes and in the peritoneal lavage of infected wild type and CCT8^−/−^ mice. During the initial infection, CCT8^T−/−^ mice had a higher frequency of ILC2 at both anatomical sites but the absolute cellularity was comparable to that of control animals (**Figure 5b, h, Appendix figure 8d**). Upon re-infection with *H. polygyrus*, the ILC2 frequency was comparable for CCT8^T−/−^ and CCT8^T+/+^ mice but the absolute cellularity was lower in mutant mice (**Figure 5h**). This finding correlated with a lack of intestinal tuft cells, thus suggesting a feed-forward loop which increases numbers of these cells (Gerbe, Sidot et al., 2016, von Moltke et al., 2016) whereby ILC2 cells do not suffice and an adaptive T cell contribution is required. This is, however, absent in the CCT8^T+/+^ mice, particularly during a secondary infection (**Figure EV 4**). In addition, the scarcity of AAMacs, eosinophils and B cells in the peritoneal lavage of CCT8^−/−^ mice demonstrated that the overall provision of type 2 cytokines in response to the nematode was inadequate to recruit an effective cellular and humoral response to *H. polygyrus*.

## Discussion

CCT captures and manipulates the folding of non-abundant intermediates from a range of substrates whereby its interaction occurs in a subunit-specific and geometry-dependent fashion (Gao, Thomas et al., 1992). We demonstrate here that a targeted loss of only the CCT8 protein compromised the function of the entire CCT complex and thus impaired the correct folding and consequent function of different substrates, including the cytoskeletal proteins actin and tubulin. Actin with its V-shaped molecular structure (Kabsch, Mannherz et al., 1990) has been identified as a prototype substrate that requires CCT’s function for its efficient folding (Gao et al., 1992), a process involving the binding to CCT4 and either CCT2 or CCT5 but spares direct contacts with CCT8 (Gao et al., 1992, Llorca, McCormack et al., 1999). The biogenesis of actin is, however, unaffected by the depletion of CCT8, a finding in line with observations in *C. elegans* where actin levels remain normal despite an impaired CCT8 function (Saegusa et al., 2014). In this experimental system aggregates including actin are not efficiently cleared from the cytoplasm causing the formation of aggregates and misfolded proteins that convey either a loss, a change or even a gain of function as a result of their toxicity.

Productive folding and processing of actin in T cells is an important prerequisite for the signaling-dependent changes in shape. Indeed, T cells adapt to different physiological conditions including, for example, the stress of the bloodstream flow, the migration to and residence in different tissues each with their bespoke microenvironments, the engagement with antigen-presenting cells and the formation of immunological synapses (Burkhardt, Carrizosa et al., 2008). Actin polymerization in T cells occurs in response to TCR-mediated activation and rapidly creates, in the presence of the Arp2/3 complex and several formins, a dynamic network of cytoplasmic and nuclear filaments. These structures play a pleiotropic role in T cell activation as they control the promotion of conjugates, the activation and nuclear import of transcription factors, and, possibly, the internalization of the TCR (Tsopoulidis et al., 2019). Conversely, inhibiting the cytoplasmic and nuclear formation of actin filaments broadly impairs T cell biology (Tsopoulidis et al., 2019).

Our results demonstrated that depleting CCT constrains the nuclear formation of actin filaments and results in a panopoly of functional changes. For instance, TCR-mediated progression of CCT8-deficient thymocytes past several intrathymic checkpoints was impaired, including the cells’ normal positive and negative selection. These findings align well with studies that have identified a role for UPR in thymocyte maturation and selection (reviewed in ref.(Kemp & Poe, 2019)). Interestingly, the observed partial block in thymocyte maturation was relatively mild despite a complete absence of CCT8 at the DP and later stages of thymocyte maturation. In stark contrast, the periphery of CCT8^T−/−^ mice is severely lymphopenic indicating peripheral T cells to be especially reliant on this complex for their maintenance and function. Indeed, both ER stress and UPR play an important role in the homeostatic maintenance of peripheral T cells and are disordered in the absence of CCT. The mass spectrometric analysis of both resting and activated CCT8^T−/−^ T cells further confirms at the protein level that several molecules involved in mitochondrial functions, including Phb1 and Phb2, are expressed at lower concentrations in the absence of CCT8. Phb1 and Phb2 form a hetero-oligomeric complex that contributes, *inter alia*, to an unfolded protein response in mitochondria (Tatsuta & Langer, 2017).

In addition to a compromised cell viability and a restricted clonal expansion, the absence of CCT in activated T cells also causes an increased ability to secrete IFN-γ and a limited efficiency to adopt a Th2 phenotype, even when exposed *in vitro* to ideal polarizing conditions. While T-bet favours the expression of IFN-γ the same factor represses the Th2 lineage commitment via a tyrosine kinase mediated interaction that disrupts the binding of GATA3 to its DNA binding motif (Szabo, Sullivan et al., 2002). We do see an upregulation of STAT1 in the CCT8^T−/−^ cells after 24 hrs of activation under non-polarizing condition. Though attractive as an explanation we could not detect T-bet or the IL-12 receptor. The upregulation of GATA3 and activation of signal transducer and activator of transcription 5 (STAT5) are two indispensable events for this differentiation process to occur.

The co-evolution of vertebrate hosts with helminths has resulted over the course of millions of years in a sophisticated immune defence that engages several effector mechanisms orchestrated by a robust Th2-type response. This reaction activates and mobilizes a suite of innate immune cells and local tissue responses (Maizels, Hewitson et al., 2012b). During the worm’s life cycle infective larvae taken up by oral route invade the mucosa of the duodenum, cross its muscle layer and reach a space beneath the serosa from where adult parasites return 8 days later to access the intestinal lumen Even though a primary infection does not resolve without drug treatment under the experimental conditions used here, a secondary exposure is completely cleared in wild-type mice when sufficient IL-4 (but not necessarily IL-13) is available to activate responsive epithelia (Anthony et al., 2006, Shea-Donohue, Sullivan et al., 2001) and AAMacs forming granulomatous cysts to encase the larvae that have penetrated the intestinal wall (Anthony et al., 2006).

The activation and expansion of B cells secreting cytokines and immunoglobulins, in particular IgG1 in a T cell-dependent fashion, are known to be critical for effective anti-*H. polygyrus* immunity (Hewitson, Filbey et al., 2015, Liu, Kreider et al., 2010, McCoy, Stoel et al., 2008). Resistance to *H. polygyrus* (re-) infection also results in the activation of group 2 innate lymphoid cells (ILC2) (Pelly, Kannan et al., 2016), which are required for the differentiation of conventional Th2 cells (Pelly et al., 2016). However, the limited capacity of CCT8^T−/−^ mice to effectively polarize their CD4^+^ T cells *in vivo* to a Th2 phenotype thwarts the clearance of *H. polygyrus* upon re-infection even in the presence of ILC2 cells. Total cellularity and the frequency of Th2-type T cells are significantly reduced in both mesenteric lymph nodes and the peritoneal lavage of infected CCT8^T−/−^ mice when compared to wild type animals. This reduction and its down-stream cellular and molecular consequences – possibly in addition with the general lymphopenia of CCT8^T−/−^ mice-explains mechanistically the animals’ inability to expel *H. polygyrus* via a robust immune response that generates sufficient IL-4. Interestingly, ILC2 in both mesenteric lymph nodes and the peritoneal lavage increase in CCT8^T−/−^ mice during the first worm exposure indicating that the initial recruitment of these cells is largely intact. However, these cells require a greatly expanded Th2 population to escalate a full-scale anti-helminth immune reaction including tuft cell proliferation and alternative activation of macrophages. Although ILC2 cells are an innate source of IL4, their reduced cellularity is overtly insufficient to create an effective immune defence in CCT8^T−/−^ mice to by-pass their defective Th2 response. This and additional findings presented here indicate that Th2-type cells are indispensable for a protective immunity against *H. polygyrus* whilst positioned upstream of AAMac, eosinophils and B cells whose activation and expansion cannot be driven by ILC2 alone.

*H. polygyrus* infections also expand and activate the host’s T_reg_ population, especially early in an infection when these cells appear to outpace the proliferation of effector T cells and modulate the immune response (Filbey, Grainger et al., 2014). An increased frequency of T_reg_-possibly as a result of low dose IL-4 exposure (Tu, Chen et al., 2017) was observed in the mesenteric lymph nodes of infected CCT8^T−/−^ animals where these cells may further inhibit Th2 immunity (Grainger, Smith et al., 2010). However, the frequency of T_reg_ was significantly reduced in the peritoneal lavage. The molecular reason for this compartmentalization of T_reg_ remains so far unknown but may reflect a deficiency in the cell’s expression of CD103 (Filbey et al., 2014) and homing to the intestine. This could result from a lack in up-regulating the gut-trophic chemokine receptor CCR9 and/or an irregular response to CCL25, a chemokine highly expressed in the epithelium of the inflamed small intestine and on postcapillary venules of the lamina propria.

In mouse strains susceptible to *H. polygyrus* infections, Th2 responses are counterbalanced by IFN-γ-producing CD4^+^ and CD8^+^ T cells (Filbey et al., 2014). The prediliction of CCT8^T−/−^ T cells to adopt a Th1 phenotype and secrete IFN-γ upon activation suppresses the formation of a type 2 immune response and inhibits cell proliferation, thus further contributing to a deviation away from protective immunity against *H. polygyrus*. It is tempting to speculate that such a shift towards a Th1-type immune response due to an absence of regular CCT8 function may be harnessed therapeutically in non-infectious, Th2-driven pathologies (e.g. allergic diseases). However, the benefit of such an imbalance towards a Th1-weighted immune reaction requires further probing, for example, in the context of an anti-viral immune response where IFN-γ discloses both anti-viral and immunomodulatory functions to combat the infection while minimizing collateral tissue damage (Lee & Ashkar, 2018).

Taken together, CCT-mediated protein folding is essential for normal T cell biology as their development, selection and function are severely harmed by a lack of normal proteostasis. Taken together, we demonstrate for the first time that the loss of CCT function in T cells disturbs normal proteostasis and the dynamic formation of nuclear actin filaments, a prerequisite for normal cell cycle progression and chromatin organisation. As a consequence, the maintenance of a normal peripheral T cell pool is severely compromised which correlates with an inadequate mitochondrial response to metabolic demands and an abnormal unfolded protein response weakening the ability to cope with activation-induced cell stress. Finally, CCT function is required for T cells to be TH2 polarized and to stage a protective immune response against *H. polygyrus* via adequate feed-forward loops engaging multiple cellular effector mechanisms.

## Methods

### Lead Contact and Materials Availability

Further information and requests for resources and reagents should be directed to and will be fulfilled by the Lead Contact, Georg Holländer.

### Experimental Model

#### Mice

The Cct8 KO mouse model (Cct8tm1a(KOMP) Wtsi, Project ID CSD45380i) was obtained from the KOMP Repository (www.komp.org) and generated by the Wellcome Trust Sanger Institute (WTSI). Targeting vectors used were generated by the Wellcome Trust Sanger Institute and the Children’s Hospital Oakland Research Institute, as part of the Knockout Mouse Project (3U01HG004080), designed to delete exon 2 in the Cct8 gene. The Cd4-Cre Transgenic mouse was originally developed at the University of Washington on C57BL/6 background (Lee, Fitzpatrick et al., 2001). Mice between 4 and 10 weeks were used for experiments. Animals were maintained under specific pathogen-free conditions and experiments were performed according to institutional and UK Home Office regulations.

#### Flow cytometry, cell sorting and cell purification by magnetic-activated cell separation

Cells from thymus, spleen, and lymph nodes were isolated from wild type and mutant mice and stained using combinations of the antibodies listed in table 1. Where needed, unmanipulated naïve CD4+ cells were enriched using magnetic separation (Milteny Biotech). Before staining, cells were resuspended at a concentration of 10×10^6^/100μL in PBS containing 2% FCS (Merck). Staining for cell surface antigens was performed for 20 minutes at 4°C in the dark. Thymocytes analysed were lineage negative, i.e. lacked the expression of CD11b, CD11c, Gr1, CD19, CD49b, F4/80, NK1.1, GL3, and Ter119. The following panels were used: Thymocyte surface panel (DN), antibodies recognizing; CD8 AF700, CD4 APC/Cy7, TCRβ Pe, CD24 FITC, CD25 efluor450, CD44 Pe/efluor610, CD69 Pe/Cy5, CD5 Pe/Cy7. Negative selection panel: CD8a AF700, CD4 FITC, TCRβ APC-Cy7, CD24 PerCP-efluor710, PD1 APC, CD25 Biotin, NK1.1 biotin, CD69 PE-Cy5, CD5 PE-Cy7, Streptavidin BV605, CCR7 PE, Helios BV421, FOXP3 PE-efluor610. Positive selection panel and SM, M1 and M2, antibodies recognizing; CCR7 BV421, CD4 Pe/efluor610, CD8 AF700, CD25 BV605, CD69 Pe-Cy5, CD5, PerCP-Cy5.5, TCRβ FITC, MHC-1 Pe, CD24 APC, Sca-1 Bv510, CD44 Pe-Cy7 Qa2 Biotin, streptavidin APC-Cy7.Treg panel: CD4 APC-Cy7, CD8a AF700, CD3e PerCP-Cy5.5, CD25 efluor450, CD69 PE-Cy5, CD5 PE-Cy7, ICOS PE, CD103 AF647, CD24 FITC, FOXP3 Pe/efluor610, CCR6 APC. Memory/naïve panel; CD4 APC-Cy7, CD8a AF700, B220 BV605, CD44 Pe/efluor610, CD62L PE-Cy7, TCRβ PE-Cy5, CD25 ef450, CD5 PerCP-Cy5.5, FR4 APC, CD73 PE. T cell polarization panels; IFNγ FITC, IL17 Pe, FoxP3 Pe/efluor610, Rorγτ APC. Where needed, staining for CCR7 was performed for 30 minutes at 37°C in a water bath, directly followed by the addition of other cell surface stains. For the identification of intracellular markers (Foxp3, IL-17, Helios, IFN-γ) Foxp3 Transcription Factor Staining Buffer Set (eBioscience) was used according to manufacturer’s instructions; intracellular stains were performed for 60 minutes at 4°C in the dark. For the identification of IL-4, cells were fixed with 2% PFA and then permeabilized with 0,1% Saponin. Prior to cytokine staining, cells were incubated with Cell Stimulation Cocktail (eBioscience) for 2 hours before adding monensin, and further incubated at 37 °C for a total of 5 hours. Cell viability was measured using LIVE/DEAD Fixable Aqua Dead Cell Stain Kit (ThermoFisher Scientific) as per the manufacturer’s instructions. After staining cells were acquired and sorted using a FACS Aria III (BD Biosciences) and analysed using FlowJo v10.

#### Detection of TNF-α production

Assay of TNF-α production was performed in accordance with (Xing, Wang et al., 2016). Briefly, total thymocytes were activated with 1 μg/mL plate bound anti-CD3 and soluble 1 μg/mL anti-CD28 in the presence of monensin (BioLegend). After 4hrs, cells were stained for surface markers (SM, M1, M2 panel), treated with Foxp3 Transcription Factor Staining Buffer Set (eBioscience) according to manufacturer’s instructions before staining for intracellular TNF-α.

#### Western blot

100, 000 DN, DP, SP4 and SP8 cells were FACS sorted from 4 old CCT8^T+/+^ (n=8) and CCT8^T−/−^ mice (n=8) in 50,000 cell batches and stored as dry cell pellets at −80°C until enough cells were collected. The pellets were lysed in 11.25 μL lysis buffer containing 25mM Tris-HCl pH8 (Merck), 50mM NaCl (Merck), 1% NP-40 (Merck), 1m DDT (Merck), 10% glycerol (Merck), and 0.2mM PMSF (Merck) in deionised water. After 15 minutes on ice, the samples were centrifuged at 15,000xg for 10 minutes at 4°C. 11.25μL of each supernatant was added to 3.75μL sample buffer containing 250mM Tris-HCl pH7 (Merck), 10% SDS (Merck), 35% glycerol (Merck) and 0.05% Bromophenol blue (Merck) and heated at 97°C for 10 minutes before gel electrophoresis. Proteins were separated on a BisTris 4-10% Gradient gel (Invitrogen) and the samples run alongside a protein standard (Biorad), before transferred to a polyvinylidene difluoride membrane (Biorad). Antibody recognizing CCT8 was added at a 1:1000 dilution, with IRDye 680RD goat anti-rabbit antibody (Licor) diluted in 1:10000 as secondary antibody, before imaging on an Odyssey imaging system (Licor). As a control for protein loading, the membrane was re-stained for GAPDH (AbCam).

#### Primary cell culture, activation, proliferation, and polarization

Naïve CD4^+^ T cells were labelled by Cell trace Violet (ThermoFisher Scientific) according to the manufacturer’s protocol and activated *in vitro* using a combination of plate bound anti-CD3 (2μg/mL) (BioLegend) and soluble anti-CD28 (2μg/mL) (BioLegend) antibodies in RPMI (Merck), containing 10% Heat Inactivated FCS (Invitrogen) and 1% Penicillin-Streptomycin (Merck). Cell proliferation was measured as dilution of the cell dye as assessed by flow cytometry.

For *in vitro* polarization, naïve T cells were freshly isolated and subsequently cultured for 4/5 days in RPMI (Merck), containing 10% Heat Inactivated FCS (Invitrogen) and 1% Penicillin-Streptomycin (Merck). Following conditions were used; Th1: 50 U/mL IL-2, 1μg/mL anti-CD28, 1μg/mL anti-CD3, 3.5ng/mL IL-12, 10μg/mL anti-IL4; Th2: 50 U/mL IL-2, 1μg/mL anti-CD28, 1μg/mL anti-CD3, 10μg/mL IL-4, 10μg/mL anti-IFNγ; Th17: 1μg/mL anti-CD28, 5ng/mL TGFβ, 10ng/mL IL-1b, 50 ng/mL IL-6, 20ng/mL IL-23, 10μg/mL anti-IFNγ, 10μg/mL anti-IL-4; Tregs: 50U/mL IL-2, 1μg/mL anti-CD28, 5ng/mL TGFβ.

#### Quantitative real-time PCR (qPCR)

cDNA was synthetized from total RNA of isolated thymocytes and T cells, and qPCR performed according to the manufacturer’s instruction (Bioline). All primers (Merck) were designed to span exon-exon boundaries and are available upon request. For the detection of *Cct8* transcripts, primers spanning exon 2 was designed. The expression of the following genes was used to assess ER stress; *Bcl2*, *Grp78*, *Atf6*, *Ire1a*, *Perk*, and *Bim*. Expression of *Gapdh* was used as an internal reference, and the delta Ct method was used in order to normalize expression. The delta delta CT (ΔΔCT) method was used for analyses of fold change, which was expressed as 2^−ΔΔCT^.

#### Phalloidin staining for nuclear actin filaments

Phalloidin staining for nuclear actin was adapted from (Tsopoulidis et al., 2019). Briefly, CD4 T cells were isolated from lymph nodes of CCT8^T+/+^ and CCT8^T−/−^ mice at 4-8 weeks of age and plated overnight (O/N) with 1μg/mL anti-CD3 and 1μg/mL anti-CD28 at 4°C at a density of 10^5^ cells/m in starvation media (RPMI (Merck), containing 0.5% Heat Inactivated FCS (Invitrogen) and 1% Penicillin-Streptomycin (Merck)) with wild type cells added CK666 (Merck), an Arp2/3inhibitor, as negative control. The next day, cells were collected in 100μL of starvation media, and allowed to adhere for 5 min on polyK-coated 35mm glass bottom dishes (Ibidi). Stimulation was performed by adding 100μL of PMA/Iono solution [1:1000) cell stimulation cocktail (Invitrogen) in starvation media] dropwise to the cell suspension for 30 sec followed by permeabilization with a 100μL mixture containing (0.3% Triton X-100 + Alexa Fluor Phalloidin 488 (1:2000) in cytoskeleton buffer [10 mM MES, 138mM KCl, 3mM MgCl, 2mM EGTA, and 0.32M sucrose (pH 7.2)] for a maximum of 1 min. Cells were then fixed with 3mL of 4% Formalin Solution (Merck) and incubated for 25 min. Fixed cells were washed twice with cytoskeleton buffer and stained with 1:500 30972 Abberior® STAR 635 (Merck) O/N at 4°C. Stained cells were washed with cytoskeleton buffer and imaged using a Leica SP8 STED microscope. To determine the number of cells with nuclear actin filaments, an average of 70 cells were scored for each sample. For the cell images used in figure 3 the background was removed using the image processing software Fiji. Further, the cell membrane and nuclear envelope were removed by segmentation using the relevant function of the software.

#### Proteomics

7575 naïve CD4 cells were isolated from lymph nodes of CCT8^T+/+^ and CCT8^T−/−^ mice at 4-8 weeks of age by FACS sorting directly into lysis buffer (RIPA buffer (ThermoFisher Scientific) + 4 % IGEPAL CA-360 (Merck)) for timepoint 0 hrs. At the same time, 10^6^ cells/mL FACS sorted, naïve CD4 T cell were activated *in vitro* using a combination of plate bound anti-CD3 (2μg/mL) (BioLegend) and soluble anti-CD28 (2μg/mL) (BioLegend) antibodies for 24 hrs before 7575 live cells were sorted directly into lysis buffer. Samples were stored at −20 °C until further processing. Samples were prepared for proteomic analysis using a modified SP3 method (Sielaff, Kuharev et al., 2017). After thawing, 25 units of Benzonase (Merck) were added and samples incubated on ice for 30 minutes. Proteins were reduced with 5 mM dithiothreitol for 30 minutes at room temperature and then alkylated with 20mM iodoacetamide for 30 minutes at room temperature. 2μL of carboxyl-modified paramagnetic beads were added to the samples (beads were prepared as in Hughes *et al*)(Hughes, Foehr et al., 2014) along with acetonitrile to a concentration of 70% (v/v). To facilitate protein binding to the beads samples were vortexed at 1,000 rpm for 18 minutes. The beads were then immobilized on a magnet, the supernatant discarded, and beads washed twice with 70% (v/v) ethanol and once with 100% acetonitrile. Washed beads were resuspended in 50mM ammonium bicarbonate containing 25ng trypsin (Promega) by brief bath sonication and incubated at 37°C O/N. After digestion, beads were resuspended by brief bath sonication and acetonitrile was added to 95% (v/v). Samples were vortexed at 1,000 rpm for 18 minutes to bind peptides, then beads were immobilized on the magnet for 2 minutes and the supernatant discarded. Beads were resuspended in 2% DMSO, immobilized on the magnet for 5 minutes and the supernatant transferred to LC-MS vials which were stored at −20°C until analysis.

Peptides were analysed by LC-MS/MS using a Dionex Ultimate 3000 UPLC coupled on-line to an Orbitrap Fusion Lumos mass spectrometer (ThermoFisher Scientific). A 75μm × 500 mm C18 EASY-Spray column with 2 μm particles (ThermoFisher Scientific) was used with a flow rate of 250nL/min. Peptides were separated with a linear gradient which increased from 2-35 % buffer B over 60 minutes (A: 5% DMSO, 0.1% formic acid in water; B: 5% DMSO, 0.1% formic acid in acetonitrile). MS1 Precursor scans were performed in the Orbitrap at 120,000 resolution using an AGC target of 4e5 and a maximum cycle time of 1 second. Precursors were selected for MS/MS using an isolation window of 1.6m/z and were fragmented using HCD at a normalized collision energy setting of 28. Fragment spectra were acquired in the ion trap using the Rapid scan rate using an AGC target of 4e3.

Raw data files from each of the independent experiments were searched separately against the Uniprot mouse database (Retrieved 2017/03/15; 59094 entries) using MaxQuant(Cox & Mann, 2008) (version, 1.6.2.6). Trypsin specificity with two missed cleavages was specified, along with carbamidomethylation of cysteine as a fixed modification, and oxidation of methionine and protein n-terminal acetylation were allowed as variable modifications. The ‘match between runs’ option was enabled. Protein quantification was performed using the MaxLFQ (Cox, Hein et al., 2014) algorithm within MaxQuant and identifications were filtered to a 1% false discovery rate on the peptide-spectral match and protein levels.

#### Transcriptomic

Two thousand naïve CD4 T cells were isolated by FACS, sorting directly into lysis buffer (RLT buffer (Qiagen)) for timepoint 0hrs. At the same time, FACS sorted, naïve CD4 T cells (10^6^ cells/mL) were activated *in vitro* using a combination of plate bound anti-CD3 (2μg/mL) (BioLegend) and soluble anti-CD28 (2μg/mL) (BioLegend) antibodies for 24 hrs before 2000 live cells were FACS sorted directly into lysis buffer (RLT buffer (Qiagen)). RNA was extracted using the RNeasy Plus Micro kit (Qiagen). RNA was processed using the RNA-Seq Poly A method and 100bp paired-end RNA-Seq was performed on the Illumina HiSeq4000 platform (Wellcome Trust Centre for Human Genetics, University of Oxford). Sequencing data was trimmed using Trimmomatic and aligned to the mm10 genome using STAR (version 2.5.3a)(Dobin, Davis et al., 2013). Reads were assigned to genes using HTSeq (intersection non-empty)(Anders, Pyl et al., 2015). Differential analysis was performed using edgeR(Robinson, McCarthy et al., 2010). Significant genes were defined as those with Benjamini-Hochberg adjusted p-values less than 0.05. ClusterProfiler was used for gene ontology analysis (Yu, Wang et al., 2012).

#### Analysis of ER stress

CCT8^T+/+^ and CCT8^T−/−^ naïve CD4 cells were activated *in vitro* using a combination of plate bound anti-CD3 (2μg/mL) and soluble anti-CD28 (2μg/mL) antibodies. As a positive control, CCT8^T+/+^ cells were also cultured in the presence of 10μg/mL Tunicamycin (Merck) to induce ER stress. After 24 and 48 hrs, viable cells were isolated by flow cytometry, RNA extracted (Qiagen), and cDNA synthesized (Bioline) for subsequent analysis by qPCR.

#### Metabolic assessments

Splenic total CD4^+^T cells were isolated by magnetic sorting using commercial kit (Miltenyi Biotec.) according to the manufacturer’s protocol. Cells were plated at a density of 1×10^6^ cells/mL and activated with Mouse T-Activator CD3/CD28 beads according to the manufacturer’s manual (ThermoFisher Scientific) in RPMI (Merck), supplemented with 10% Heat Inactivated FCS (Invitrogen) and 1% Penicillin-Streptomycin (Merck) for 24hrs. For all the following analysis, live cell stains were performed *in situ* followed by incubation in the dark, in a humidified gassed (5% CO2) incubator at 37°C for 30 min. Active mitochondria were measured by adding 0.5ng/mL Mitotracker DR (Invitrogen), while MitoID (Enzo Life Sciences) was added to measure total mitochondria, according to the manufacturer’s protocol. Reactive oxygen species was measured by adding 5μM MitoSox (ThermoFisher Scientific) to the cell cultures. Lipid droplets were analysed by the addition of 1μg/mL Nile red, for measurement of palmitate uptake cells were incubated with 1μg/mL BODIPY-FLC16 (ThermoFisher Scientific) and for uptake of glucose 5μg/mL 6-NBDG (Invitrogen) was added. Apoptotic cells were detected using an Annexin-V/7AAD kit (Biolegend) with or without the addition of 1μg/L N-Acetyl-L-cysteine (Merck). Mitochondrial membrane potential (Δψm) was measured with 2μM JC-1 dye (ThermoFisher Scientific) by flow cytometry according to the manufacturer’s directions. All staining experiments were performed at least three times with biological replicates.

#### Measurement of metabolic flux

Cellular metabolism was measured using an XF96 cellular flux analyser instrument from Seahorse Bioscience. OXPHOS was measured using a Mitostress test kit (Agilent Technologies) according to the manufacturer’s instructions. Primary T cells, 3 × 10^5^ per well, were cultured in RPMI with no sodium bicarbonate and 1% FCS, 20mM glucose, 2mM pyruvate, and 50μM β mercaptoethanol (pH 7.4) at 37°C for these assays. Glycolysis was measured via ECAR measurements from the mitostress test data. Final drug concentrations used in the Seahorse assays were; Oligomycin 1μM, FCCP 1.5μM, Rotenone and Antimycin A both at 1μM. Primary data was analysed using Wave desktop software from Agilent Technologies.

#### Imaging Flow cytometry

Metabolic function was assessed by a 2 camera, 12 channel ImageStream X MkII (Amnis Corporation) with the 60X Multimag objective and the extended depth of field option providing a resolution of 0.3 μm per pixel and 16 μm depth of field. Fluorescent excitation lasers and powers used were 405 nm (50 mW), 488 nm (100 mW), and 643 nm (100 mW). The side scatter laser was turned off to allow channel 6 to be used for PE-Cy7. Bright field images were captured on channels 1 and 9 (automatic power setting). A minimum of 30,000 images were acquired per sample using INSPIRE 200 software (Amnis Corporation) and analyzed by the IDEAS v 6.2 software (Amnis Corporation). A colour compensation matrix was generated for all 10 fluorescence channels using samples stained with single color reagents or antibody conjugate coated compensation beads, run with the INSPIRE compensation settings, and analyzed with the IDEAS compensation wizard. Images were gated for focus (using the Gradient RMS feature) on both bright field channels (1 and 9) followed by selecting for singlet cells (DNA intensity/aspect ratio) and live cells at the time of staining, i.e., Live/Dead aqua low intensity (channel 8) or low bright field contrast (channel 1).

#### Infection with *Heligmosomoides polygyrus*

CCT8^T+/+^ and CCT8^T−/−^ mice were infected with 200 L3 *H. polygyrus* larvae by oral gavage. Adult parasites were eliminated by 100mg/kg of pyrantal embonate (PyratapeP) by oral gavage. For secondary infection, drug-treated mice were challenged with another 200 L3 *H. Polygyrus* larvae by oral gavage 10 days later. Faecal egg counts were used to assess the parasite burden and efficacy of the anti-helminthic treatment throughout the experiment and adult worm burdens were determined using standard procedures (Camberis, Le Gros et al., 2003).

#### IgG1 serum ELISA

ELISA plates were coated with *H. polygyrus* excretory/secretory product (HES) (Johnston, Robertson et al., 2015) at 1μg/ml in PBS O/N at 4°^C^, washed 5 times with PBS containing 0.1% Tween (PBS-T) and then blocked using 2% BSA in PBS for 2 hours at 37°^C^. The plates were then washed a further 5 times in PBS-T, followed by mouse serum added at a 1:500,000 dilution in blocking solution in the first well with serial 3-fold dilutions across the ELISA plate and left to incubate in the fridge at 4°C O/N. The following day the plates were washed with PBS-T and incubated with goat anti-mouse IgG1-HRP (Southern Biotech, USA) at 1:6000 dilution in blocking buffer and left for 1 hour at 37°^C^. After incubation the plates were washed 5 times in PBS-T and a further 2 times in distilled water, after which ABTS substrate (KPL, USA) was added and the plates were read at 405nm after 2.5 hours, using PHERAstar FS plate reader (BMG LABTECH, Germany).

#### Tuft cell staining

Small intestines were isolated from mice shortly after euthanization and flushed with PBS, the gut was then inverted onto a wooden skewer and placed in 10% neutral buffered formalin (NBF, Merck) solution for 4 hours. The tissue was sliced longitudinally to remove from the skewer and tissue was then rolled up beginning at the ileum using the Swiss roll method as previously published (Bialkowska, Ghaleb et al., 2016). The tissue was stored in NBF solution O/N then placed into 70% ethanol prior to paraffin embedding. Thin sections of paraffin-embedded tissue were de-waxed and stained with anti-DCAMLK1 at 1:1000 (Abcam, UK) and secondary stained using anti-rabbit-FITC, control slides were incubated with rabbit IgG (Abcam, UK). Once stained slides were mounted in DAPI mounting media (Vector Laboratories, UK) and imaged using the 10x objective on a Leica DiM8 microscope (Leica, Germany); images were analysed using ImageJ/Fiji.

### Quantification and Statistical Analysis

#### Statistics

Standard deviations and P-values were determined using GraphPad Prism software (Graph Pad Software Inc). P-values were calculated using a two-tailed unpaired Student’s t-test with 95% confidence interval. Each figure legend indicates methods of comparison and corrections.

## Data Availability

RNAseq data are available at the NCBI Gene Expression Omnibus under accession number GEO: GSE147209.

The mass spectrometry proteomics data have been deposited to the ProteomeXchange Consortium via the PRIDE partner repository with the dataset identifier PXD018834, Reviewer account details:

Username:
Password: FBZnFVxe

## Acknowledgements

This work was supported by the Wellcome Trust through an Investigator Award to RMM (Ref. 106122), and the Wellcome Trust core-funded Wellcome Centre for Integrative Parasitology at the University of Glasgow (Ref: 104111). BEO was supported by the Norwegian Research Council (Ref. 250030). SD was supported by a NDM studentship award (Oxford, UK).

## Author Contributions

Conceptualisation: G.A.H., and N.T.; Methodology: G.A.H., and N.T; Software: S.D., A.H., Investigations: B.E.O., S.M., N.P., S.D., A.H., I.A.R., E.S., D.H., M.P.J.W., R.M.M. and M.E.D.; Resources: G.A.H., D.H., R.M.M., R.F., B.M.K.; Data Curating: A.H., S.D.; Writing: G.A.H, B.E.O., S.M. and A.H. with contributions from all.

**Expanded View Figure 1**

Selection and maturation of cortical and medullary thymocytes in the absence of CCT8 expression. Data from CCT8^T+/+^ and CCT8^T−/−^ mice are shown in grey and white bars, respectively. (**a**) Targeting strategy for the conditional deletion of exon 3 of the *Cct8* locus. FRT: flippase recognition target; loxP: locus of X-over P1; Flp: Flipase; Cre: Cre-rcompinase. (**b**) *Cct8* transcripts in CCT8^T+/+^ and CCT8^T−/−^ thymocytes at indicated stages of development. qPCR analysis normalised to GPDH and displayed as 2^−ΔΔCt^ CT values. (**c**) Differentiation of linage negative CD24^−^TCRβ^−^ CD4^−^CD8^−^ thymocytes. Lineage negativity is defined as the absence of CD11b, CD11c, Gr1, CD19, CD49b, F4/80, NK1.1, GL3, and Ter119 expression. Left: contour plots and gating, see also Figure EV 2. (**d**) SPCD4 thymocyte maturation. Left: Contour plots showing gating. Right: (i) Individual maturational stages, qualified as immature (CD24^+^CCR7^−^TCRβ^+^: S1), semimature (CD24^+^CCR7^+^TCRβ^+^: S2) and mature (CD24^−^CD69^−^ TCRβ^+^: S3). (ii) Maturational progression between sequential developmental stages. (**e**) SPCD8 thymocyte maturation. Left: Contour plots showing gating. Right: (i) Individual maturational stages, qualified as immature (CCR7^+^TCRβ^+^: S1), semimature (CD24^+^CD69^+^: S2) or mature (CD24^−^CD69^−^: S3). (ii) (ii) Maturational progression between sequential developmental stages. *p<0.05, **p<0.01, ***p<0.001, ****p<0.0001. Data in panel b-e were calculated by Student’s t-test. Bar graphs show the mean ±SD and are representative of one of 2 independent experiments with 3 samples each.

**Expanded View Figure 2**

Proteomic analysis of activated CD4^+^ cells. Bar graphs display data from CCT8^T+/+^ and CCT8^T−/−^ mice in grey and white bars, respectively. (**a**). Percentage of live (i.e. DAPI^−^) cells after *in vitro* stimulation for the indicated length of time in the presence or absence of N-acetyl-L-cysteine (NAC: 5mM). (**b**) Go-analysis of differentially expressed genes in (i) CCT8^T+/+^ and (ii) CCT8^T−/−^ CD4^+^ T cells stimulated *in vitro* for 24h by CD3 and CD28 crosslinking. (**c**) Superoxide mediated oxidation of MitoSOX™. Cells were stimulation for 20h by CD3 and CD28 crosslinking. Representative dot plot analysis (left) of data shown in bar graphs (right). (**d**) Ratio of mitochondrial membrane potential (as assessed by mitoTracker) and mitochondrial mass (as measured by MitoID-red specifically bound to cardiolipin in the inner mitochondrial membrane) in (i) CD62L^+^CD44^−^ and (ii) CD62L-CD44+ CD4+ T cells stimulated *in vitro* for 20h by CD3 and CD28 crosslinking. (**e**) Mitochondrial membrane potential assessed by J-aggregate forming lipophilic cation 5,5′,6,6′-tetrachloro-1,1′,3,3′-tetraethylbenzimidazolcarbocyanine iodide (JC-1) in CD4^+^ T cells stimulated *in vitro* for 20h by CD3 and CD28 crosslinking. (**f**) Volcano plots of RNA transcripts in wild type (right) vs. CCT8^T−/−^ left CD4+ T cells (i) before (0h) and (ii) after (24h) *in vitro* stimulation by CD3 and CD28 crosslinking. (**g**) Correlation of transcriptomic (x-axis) and proteomic (y-axis) ratio data in CD4^+^ T cells after *in vitro* stimulation for 24h by CD3 and CD28 crosslinking. Identical transcript/protein entities are shown in red. Panels c and e show on the left representative contour plots and gating. *p<0.05, **p<0.01, ***p<0.001, ****p<0.0001. Data in panel a was calculated by Student’s t test adjusted for multiple comparisons, data in panels b-d were calculated by Student’s t test. Bar graphs show the mean ±SD representative of 2 independent experiments with 3 (c-d) and 4 samples each (panel a).

**Expanded View Figure 3**

Peripheral T cell function in the absence of CCT8 expression. (**a**) Schematic of unfolded protein response (UPR) in response to unfolded or misfolded proteins. Components whose transcripts were examined are shown in bold. (**b**) *In vitro* Th17 polarization of separately labelled CCT8^T+/+^ and CCT8^T−/−^ CD4^+^ T cells stimulated in co-cultures. Left bars show the frequency of Th17 cells in cultures where CCT8^T−/−^ T cells were labeled with cell trace violet (CVT) labelled; right bars show the frequency of Th17 cells in cultures where CCT8^T+/+^ T cells were labeled with CVT. (**c**) Determination by Seahorse XF Analyser of (i) basal oxidative phosphorylation, (ii) spare respiratory capacity, and (iii) basal extracellular acidification rate (ECAR) in CCT8^T+/+^ and CCT8^T−/−^ CD4^+^ T cells stimulated for 20h by CD3 and CD28 crosslinking. OCR: oxygen consumption rate. *p<0.05, **p<0.01. Data in panel b was calculated by Student’s t test, and data in panel c was calculated by Student’s t test adjusted for multiple comparisons. Bar graphs show the mean ±SD representative of two independent experiments with three replicates (panel b) and display the combined results of three independent experiments with 14 CCT8^T+/+^ samples and 23 CCT8^T−/−^ samples (panel c).

**Expanded View Figure 4**

The response to *H. polygyrus* infection in CCT8-deficient mice. (**a**) Frequency of B cells in peritoneal lavage (i) and the mesenteric lymph nodes (ii) of CCT8^T+/+^ (grey bars) and CCT8^T−/−^ mice (white bars). (**b**) Tuft cells in the small intestine of *H. polygyrus* infected CCT8^T+/+^ and CCT8^T−/−^ mice 14 days after primary and re-infection as indicated. (i) Sections shown represent data from mice with median egg counts in each experimental group. For Dclk1 staining: yellow; DAPI: blue. Scale bar in right lower corner represents 20 μm. (ii) Number of tuft cells in wild type (grey bars) and mutant (white bars) mice. *p<0.05, **p<0.01, ***p<0.001, ****p<0.0001. Data was calculated by Student’s t test adjusted for multiple comparison. Bar graphs in panel a show the mean ±SD representative of one of 2 independent experiments with at least 4 samples (CCT8T+/+ primary infection: 5 mice, secondary infection 5 mice; CCT8T−/− primary infection 4 mice, secondary infection 6 mice). Data in panel b is representative of 4 separately analysed sections per experimental group.

